# Quantum Calculations Of A Large Section Of The Voltage Sensing Domain Of The K_v_1.2 Channel Show That Proton Transfer, Not S4 Motion, Provides The Gating Current

**DOI:** 10.1101/154070

**Authors:** Alisher M. Kariev, Michael E. Green

## Abstract

Quantum calculations on much of the voltage sensing domain (VSD) of the K_v_1.2 potassium channel (pdb: 3Lut) have been carried out on a 904 atom subset of the VSD, plus 24 water molecules (total, 976 atoms). Those side chains that point away from the center of the VSD were truncated; in all calculations, S1,S2,S3 end atoms were fixed; in some calculations, S4 end atoms were also fixed, while in other calculations they were free. After optimization at Hartree-Fock level, single point calculations of energy were carried out using DFT (B3LYP/6-31G**), allowing accurate energies of different cases to be compared. Open conformations (*i.e.,* zero or positive membrane potentials) are consistent with the known X-ray structure of the open state when the salt bridges in the VSD are not ionized (H^+^ on the acid), whether S4 end atoms were fixed or free (closer fixed than free). Based on these calculations, the backbone of the S4 segment, free or not, moves no more than 2.5 Å upon switching from positive to negative membrane potential, and the movement is in the wrong direction for closing the channel. This leaves H^+^ motion as the principal component of gating current. Groups of 3-5 side chains are important for proton transport, based on the calculations. Our calculations point to a pair of steps in which a proton transfers from a tyrosine, Y266, through arginine (R300), to a glutamate (E183); this would account for approximately 20-25% of the gating charge. The calculated charges on each arginine and glutamate are appreciably less than one. Groupings of five amino acids appear to exchange a proton; the group is bounded by the conserved aromatic F233. Dipole rotations appear to also contribute. Alternate interpretations of experiments usually understood in terms of the standard model are shown to be plausible.

## INTRODUCTION

The mechanism of gating of voltage gated ion channels has been the subject of study ever since their existence was proposed by Hodgkin and Huxley^1^. They pointed out that the response to depolarization of the membrane would have to be preceded by a capacitative current, called gating current, as charge rearranged in response to the changing electric field. This was first measured in 1974 by two groups, Keynes and Rojas^2^, and Armstrong and Bezanilla^3^. The channel structure turned out to be tetrameric, exactly so for K^+^ channels, not exactly for Na^+^ channels; in this paper we will be concerned with a K^+^ channel. Each channel contains four domains, each with a voltage sensing domain (VSD). Each VSD is composed of four transmembrane (TM) segments; each domain also has 2 TM segments that contribute to the 8 TM segments forming the pore through which the ion traverses the membrane. The way in which the VSDs bring about the opening of the pore as the membrane depolarizes has long been an active field of investigation. Horn and coworkers adapted the substituted cysteine accessibility method (SCAM)^4^ to the VSD; the VSD TM segment attached, via a linker, to the pore segments, S4, has arginines (and usually one lysine, typically near the intracellular end of the S4 segment) in every third position; for ion channels, the SCAM method was originally applied, by Horn and coworkers, to R➔C mutations in that segment, plus reaction of the cysteine with methanethiosulfonate (MTS) reagents that kill the channel; the method assumes that the cysteine must reach the surface of the membrane to react, because the cysteine must ionize in order to react, with ionization considered impossible in the putatively low dielectric constant medium in the interior of the membrane. By determining whether the membrane reacted when the MTS reagent was added to the intracellular or extracellular side, or both, or neither, of the membrane, Horn and coworkers concluded that the cysteine, hence the entire S4 segment, had moved toward the intracellular side of the membrane when polarized, the extracellular side when depolarized^5^. The latter would open the channel by pulling on the linker between the S4 segment and the pore S5 segment. X-ray determination of the K^+^ ion channel structure by MacKinnon and coworkers^6^ (first KcsA, a channel without a VSD, but then the open state of channels with VSD) made possible a specific discussion of mechanism. Only the open state was determined, because the channel could only be crystallized in the absence of the field. The extent of the displacement of the VSD has been the subject of extensive research since, and the SCAM method has been supplemented by other experiments, especially fluorescence quenching, plus molecular dynamics (MD), all of which have been interpreted in terms of what has become the standard paradigm. This may be summarized as: 1) membrane polarized: the S4 TM segment of the VSD is “down” (intracellular), pushing, through the S4-S5 linker (the S5 TM segment being part of the pore), to close the intracellular gate; 2) membrane depolarized: the S4 rises (moves extracellularly), pulls on the S4-S5 linker, separating the residues at the gate, making it open for the ions to pass through the pore, giving the observed ion current. There have been a number of proposals for the details of the motion involved, which are not necessarily consistent with each other. We avoid discussing these details, in favor of presenting an alternative that does not move the S4 to either surface. The SCAM assumption that the cysteine must move to a membrane surface to react is not necessarily correct in this instance. The cysteine side chain is essentially a single sulfur atom when ionized. If it replaces an arginine, as in the original experiments, there is a large cavity; the arginine side chain, guanidinium, is much larger than a single atom; consequences are discussed below. The cysteine can ionize in the cavity that is left by the mutation, between the outer and inner membrane surfaces, without the backbone moving to the surface. The standard models all require that the arginines remain charged throughout the entire gating cycle, that they exchange partners with alternate charged glutamates or aspartates, thus limiting the energy cost of breaking the salt bridge, so that there is limited activation energy cost along the way. However, a Q_10_ value (increase in gating probability with 10°C rise in temperature) of 1.5, (which is roughly in the neighborhood of values for some channels^7^) corresponds to about 30 kJ for gating current. The literature suggests that there may be non-Arrhenius behavior in gating, however, so this may not be a well-defined value. The H_v_1 channel, which resembles a VSD through much of the transmembrane sections, shows this behavior^8^ too. Since this channel transmits protons, it shows that protons can traverse the VSD. Another set of experiments, in which the end arginines are mutated to histidine, allow the VSD to transmit protons as well^9^. Our calculations show that proton transfers do occur under the influence of the field, leaving the arginines not always charged, and never completely charged, so that the fundamental assumption of the standard model is called into question. In addition, the S4 segment does not move more than about 2.5 Å relative to the S1,S2,S3 block, and not in the direction expected with standard models, so that even if the arginines remain ionized, they do not contribute to gating current.

Green long ago proposed an alternate gating mechanism in which the gating current was the motion of protons through the VSD, rather than the motion of positively charged arginines ^10^. Kariev and Green have discussed a possible way to understand the evidence supporting the standard paradigm that is compatible with the model we propose. The effects of R➔C mutation can be understood by looking at the difference in size of cysteine and arginine ^11^. There will be enough space that the cys could transfer a proton to another amino acid, as there are acid residues that were originally in salt bridges with the arginine, and would be missing their partner in the cys mutant; this would account for the required ionization of the cys. A fuller discussion of the reinterpretation of the evidence for the standard models is given in previous work^11^. Here we show a step in which proton motion can contribute to gating current, while the S4 backbone remains stationary.

The TASK 2-pore channel is a potassium channel somewhat analogous to the K_v_1.2 channel, but pH gated. Niemeyer et al found a single arginine residue was responsible for the pH sensitivity ^12^; the pK of the relevant arginine had shifted to 8.0, over four units. They proposed that this was a consequence of the hydrophobic environment of the arginine. While we do not find the environment of all the arginines in the VSD to be so hydrophobic, the fact that the gating of a K^+^ channel depends upon a large pK shift is in accord with our proposal; the environment is different for different arginines, but a similar effect is possible where the proton needs to transit. The protonation states in our model are necessarily transient. (Also, recall that standard models use SCAM, including the assumption of a hydrophobic environment for S4, as the reason the S4 must move to the membrane surface to react.), In addition to the arginines and glutamates, other residues, including a tyrosine, play an important role in the part we have calculated so far. Calculations of the structures and energies of the VSD with H^+^ in several possible positions leads to a possible H^+^ path through the VSD, which includes changes in ionization states of several residues, and groupings of three to five amino acids that participate in H^+^ transfer. The path appears to be modulated by a phenylalanine which is well conserved. As this phenylalanine is near the edge of the section that was calculated, further work will be needed to determine exactly how it plays its role.

Several other matters appear to be important for the gating mechanism:

i. One involves the sharing/delocalization of a proton. In earlier work, we have examined, by use of quantum calculations, the effects of water with salt bridges ^13^. This showed, in much smaller systems, that a proton could be part of a ring of hydrogen bonds that included water, which in turn affected the state of ionization. It is even possible to have what is essentially a resonance hybrid of hydrogen bonded structures.
ii. Second, an arginine, if it moves into the headgroup region of the membrane, will be held in a complex with the phosphates of the membrane lipids. This makes optimizations with S4 fixed more appropriate than with S4 free; we consider this further in Results, section b, and Discussion, section c.
iii. We have also examined the way the pore responds to opening; this requires that an incoming K^+^ be held, in effect complexed at the entrance to the pore, to allow the incoming ion to push an ion in the cavity of the pore toward the selectivity filter, rather than have that ion push the incoming ion back out ^14^. It appears that the location where the ion is held may be at the top of the T1 moiety that hangs below the membrane, below the PVPV conserved sequence at the membrane surface.
iv. We have also considered the possibility of proton tunneling ^15^, which would correspond to the “piquito”, the rapid rise time initial component of gating current ^16^; to account for the piquito it has been proposed that there is discontinuous movement of side chains in a complex energy landscape^17^. However, the tunneling proposal provides a mechanism that directly leads to the observation. Such an initial step in the gating current also suggests the crossing of a threshold as the membrane depolarizes, allowing the energy levels of the initial and final proton positions to match, as needed to allow tunneling. If there are multiple channels, with the threshold energy (potential) distributed over channels with a width in energy of about thermal energy (k_B_T, Boltzmann’s constant x temperature), one can reproduce the observed current voltage curve as well as the “Boltzmann” curve does ^18^.

Experimental indications that proton transport in the VSD should be considered include:

i. It is known that substituting histidine for one of the end arginines allows a proton current to pass completely through the VSD ^9, 19^. This tells us that proton passage is possible; it only remains to work out the path.
ii. The H_v_1 channel, which transmits protons, has a structure very similar to that of the VSD in one section of the channel, and with a structure that follows a similar pattern in the remainder, except for a section in which the protons appear to be redirected, toward the pore for the VSD and to the membrane surface for H_v_1. A recent review by deCoursey et al ^20^ discusses the evidence pointing to the proton path; they conclude that there is a small shift of S4 (“one click”), corresponding to a single exchange of partners, to produce gating. However, their evidence could be understood even without this shift. H_3_O^+^ may also be involved^21^.
iii. Zhao and Blunck have recently shown that a truncated VSD forms a cation channel with a strong preference for protons ^22^. While not a surprise in itself, and not entirely incompatible with standard models, this finding adds to the evidence that protons can, and should, move through the VSD.
iv. Proton transfer has also been reported in other membrane proteins, including cytochrome c (cyt c)^23^.

The majority of our path is not a water wire. It is necessary to determine how the H^+^ moves, but there is already sufficient evidence to show that it does move (there may be a need to explain separately how certain R➔H mutants turn the VSD into a complete H^+^ channel, but for purposes of understanding how the VSD transmits protons internally (capacitative gating current), it is not necessary to understand how the H^+^ passes the intracellular and extracellular ends of the VSD.

There have been extensive MD studies on VSDs; only a sample of recent studies is referred to here ^24^. A very recent example uses the Drude polarizable potential on a model that more nearly resembles a Na^+^ channel^25^. These papers may also include experimental data. However, classical force fields cannot usually describe proton transfer, nor do they get hydrogen bonds consistently correct, when different length hydrogen bonds are involved. The new Sun and Gong paper with Drude potentials proposes a partial trajectory for S4, but does not agree with other MD simulations. By including polarizability, it should improve its accuracy to some extent, but it still could not allow for proton transfer, hence cannot speak to the question of whether it exists. No MD has attempted to account for experimental observations of proton transfer. There is another problem with MD calculations: gating transitions must eventually return the channel to its original state, albeit not necessarily by the same path; there is evidence of hysteresis^26^, so there must be alternate paths. One expects channels to live for tens of thousands of cycles of opening and closing before the channel protein is replaced (e.g., if there is a ten minute minimum lifetime, and the channel opens at about 10 Hz on average—and some channels operate more rapidly than this—with 4 domains, this means 24,000 refoldings). This means that whatever transitions are made in gating, it must be possible to return with essential certainty to the original. The standard models, with S4 moving more than 10 Å, effectively unfold the protein, or at last a substantial section of it. Refolding with the high accuracy required, without the protein being trapped in a local minimum in any cycle, seems very difficult. Known membrane protein refolding appears to require chaperonins or equivalent ^27^. Ion channels like K_v_1.2 do not appear to have any long-term chaperonins; an initial insertion chaperonin would not help here, if it existed. At this point this is only a plausibility argument. However, the idea of a large scale unfolding/refolding event to open the channel does look somewhat implausible; the versions that have been proposed require major unfolding of the protein, internally, not at hinges as is the case with the great majority of enzymes. This would have to be followed repeatedly by refolding, possibly along a different path, without the protein becoming trapped in a local energy minimum. So far, we have not seen MD simulations, even with a polarizable potential, that show that they can return the protein reliably, in multiple repeats, to the original state.

Here we present the results of quantum calculations on larger sections of the VSD than any previously reported. These calculations are consistent with the idea that proton motion constitutes the gating current. We propose a specific path for part of this motion, based directly on the calculation, with determination of the energies of the states of the VSD with the proton in the proposed sequence of positions. The conformation of the VSD is optimized in the quantum calculation, and negligible motion of arginines is found, but the proton motion can account for the gating current; the motion of the S4 backbone is of the order of 2 Å, and not entirely in the correct direction for it to contribute to gating current. Furthermore, the conventional attribution of the ionization states of the salt bridges leads, in our calculations, to positions of the salt bridge components that differ seriously with the X-ray structure at 0 V; however, switching proton positions, hence ionization states, from ionized to neutral salt bridges, leads to interatomic distances that are a good match for those determined by X-ray structures. Since the X-ray structures do not show the protons, probably the most accurate way to get these ionization states is to use quantum calculations. We determine whether the energies of the proton-transferred cases are lower or higher than the energies of the corresponding ionized salt bridge structures, which depends in part on the applied external field. Furthermore, the structures that transfer protons often must be considered in groups of three to five residues; one member in a group we have calculated is a phenol, tyrosine. A millisecond rate of proton exchange for a tyrosine has been shown to be possible by Takeda and coworkers^28^. These structures appear in the results of the calculations for the upper (extracellular) part of the VSD. It is possible to postulate a complete path for the H^+^ using residues intracellular to the calculated residues; but these remain to be calculated. Three calculated cases are shown in Fig. 1, and in a different form, Fig. 2.

**Fig. 1:**
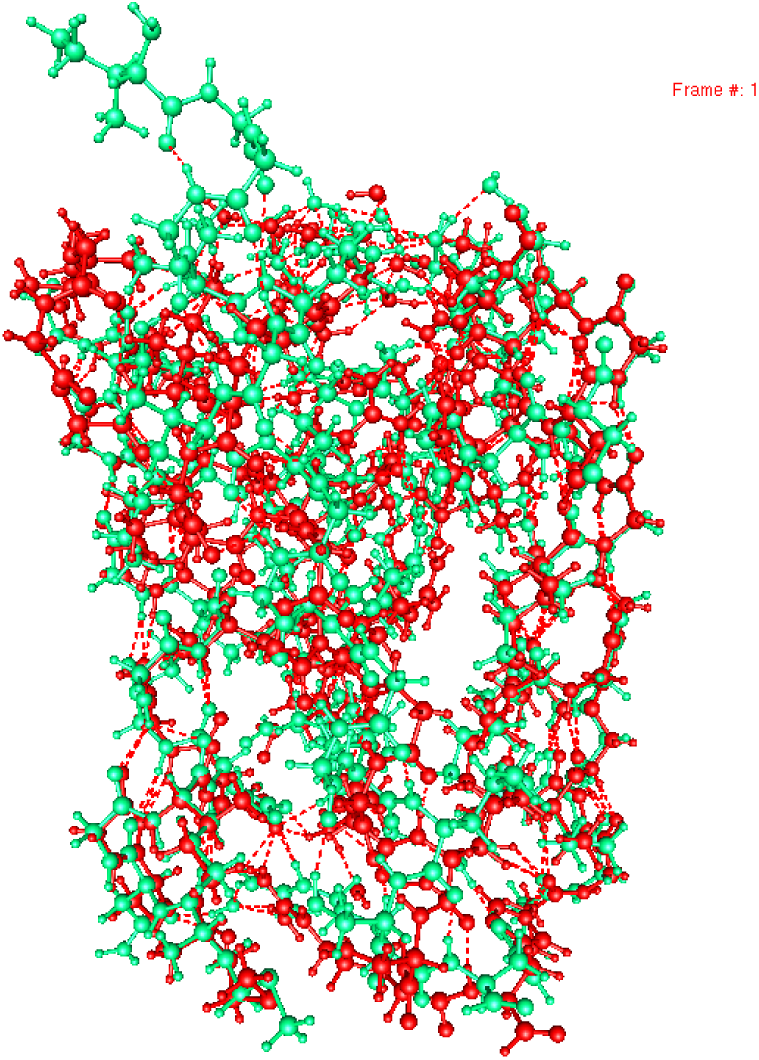
Comparison of X-ray structure (red) and optimized V<0 structure (green). Optimized structure differs from X-ray structure by ≈1 Ắ, generally less (see Table 3). The only exception is I292, which, absent other extracellular atoms, pops into open space. S4 shows no intracellular motion at all.

**Fig. 2:**
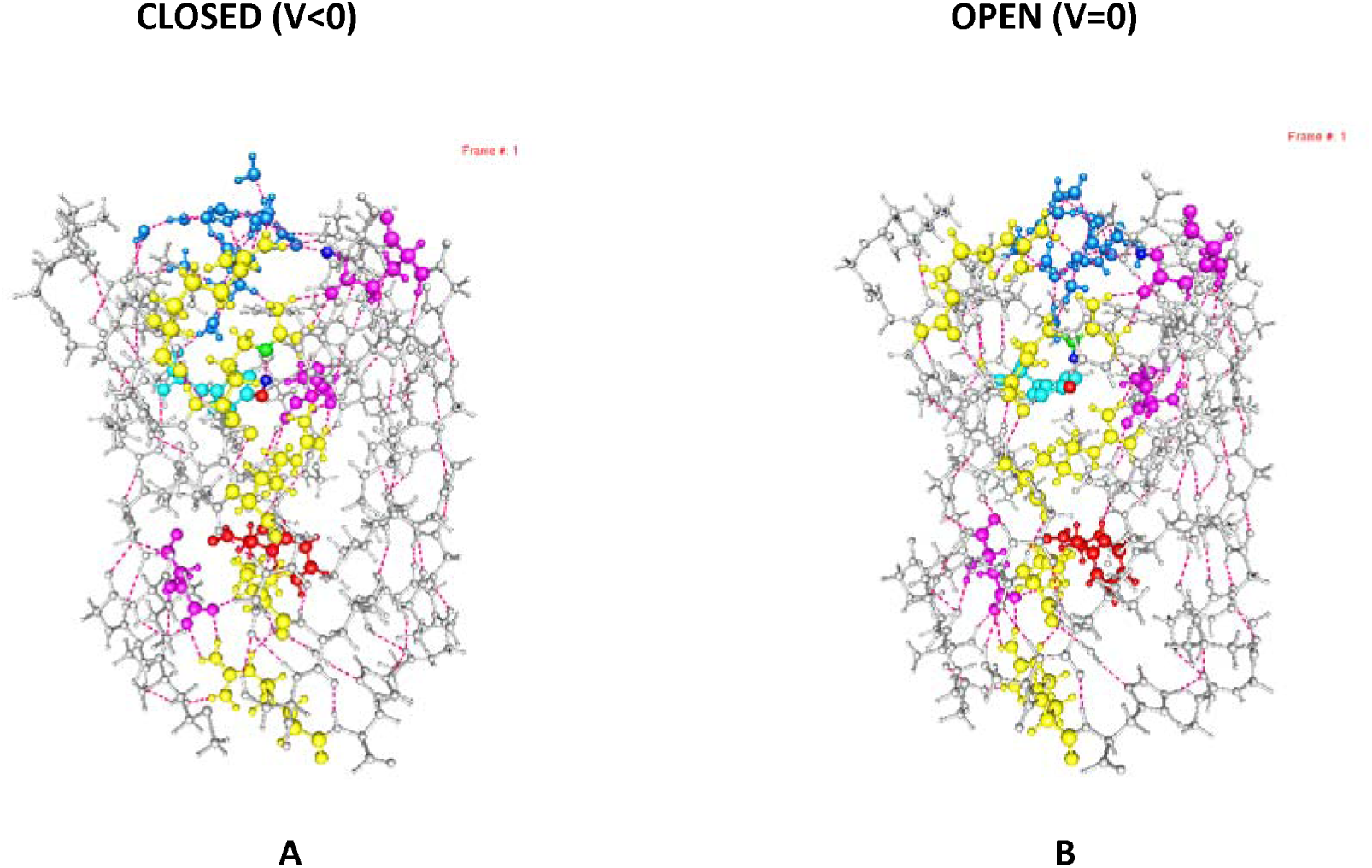
Optimized structures, S4 fixed: A) E183 protonated, V<0 (closed) B) E183 protonated, V=0 (open); **A)** proton neutralizing E183 from R300, which is neutral, while Y266 is uncharged; **B)** the proton comes from Y266, which becomes an anion, and reaches E183 by way of R300, which remains positively charged. The proton giving R300 charge resides on nitrogen NE, not on the NH_2_ amines of the guanidinium. Colors: yellow, arginine; magenta, acids; red, F233; light blue, Y266, deep blue, water. with end atoms of S1/S2/S3/S4 all fixed. The energy values, given in Table 4 (see also Fig. 3), show these to be the minimum energy cases in the open and closed paths. There is appreciable rotation of F233 between open and closed cases, and the relative arrangements of Y266 (pale blue) and E183 also change between open and closed cases.

## CALCULATIONS

We have carried out optimizations (i.e., energy minimizations) with 976 atoms (24 water molecules, 904 protein atoms from 43 amino acids, 3700 electrons, and 8,178 basis functions), varying four parameters: *i)* field; *ii)* S4 free/fixed; *iii)* proton in the salt bridge involving R297 shifted, and in one case with a proton shifted from R300; *iv*) initial direction of the average dipole of the water cluster (this turned out to make relatively little difference). In each calculation, 42 atoms from S1, S2, and S3 are fixed, all from the ends of the segments. In the *S4 free* case these were the only fixed atoms. In *S4 fixed* cases, 19 atoms from the ends of the S4 segment were also frozen, so that S4 could not move vertically. While it might seem that this defeats the purpose of the computation, comparison of S4 free, S4 fixed, and the X-ray structure shows that the S4 fixed open state reproduced the local X-ray structure, while S4 free was actually less accurate. Furthermore, even when S4 was free, it failed to move in a manner that could produce gating current. We will discuss this below, but it appears that S4 fixed is the system that comes closest to reproducing the actual condition of the system. Water molecules were always free. These calculations tell us whether it made a difference to the final positions of the S4 backbone if the backbone end atoms were fixed, or whether the interactions with S1, S2, and S3 were sufficient to keep the S4 in place, or nearly so. One additional case could be of interest: S4 free and the field set at +10^7^ Vm^-1^, equivalent to +70 mV across the membrane. (For all calculations, 10^7^ V m^-1^ is considered equivalent to 70 mV across the channel; in all cases, the field is applied as 10^7^ V m^-1^.) This should drag S4 in the opposite direction, not only open but pulled even further outward, to see whether in fact S4 could respond to such a field, or whether it would be trapped in its normal position. This large a positive field is on the edge of what is possible in many experimental preparations; still, the S4 backbone moved <2.5 Å.

All optimizations were done at HF/6-31G* level, using Gaussian09 (versions D or E) ^29^. The 976 atom calculations included the full amino acids shown in Table 1

**TABLE 1.**
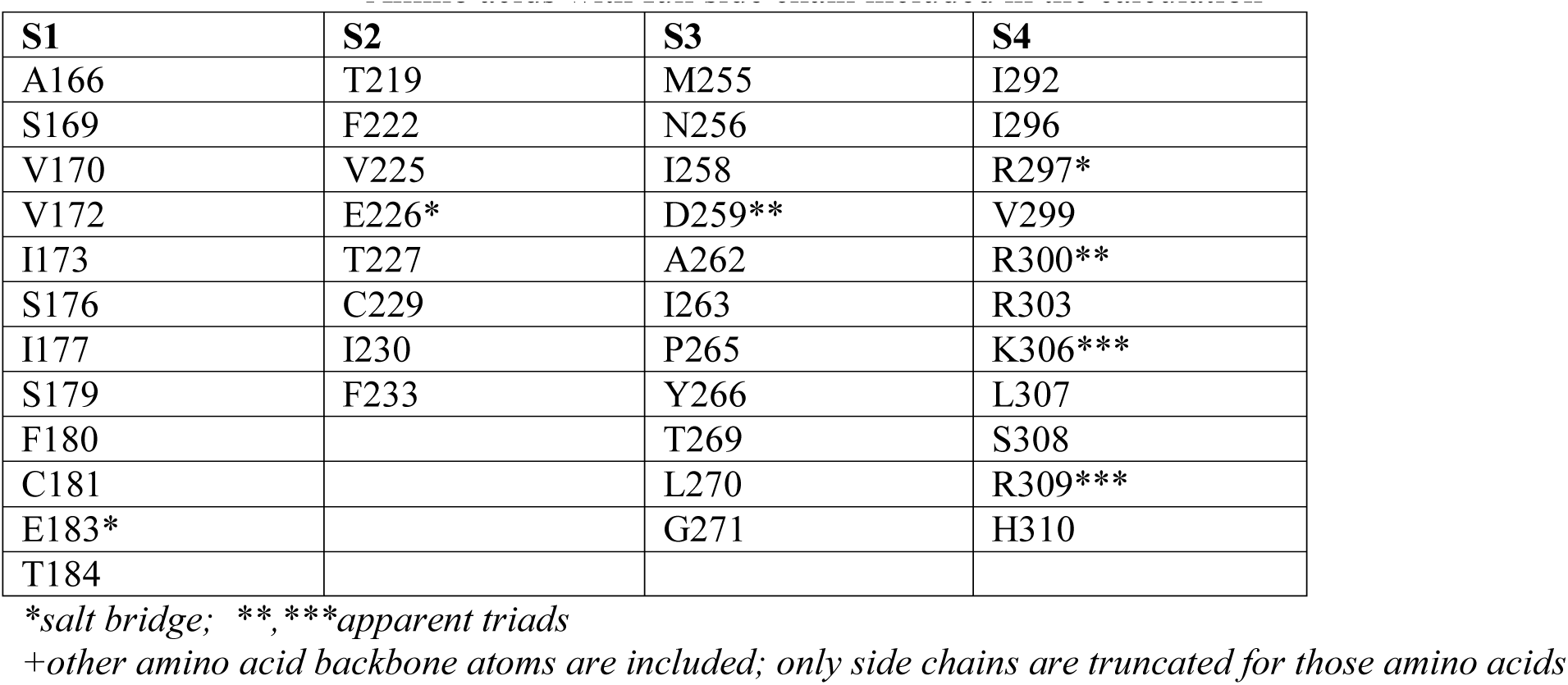
Amino acids with full side chain included in the calculation^+^

| S1 | S2 | S3 | S4 |
| --- | --- | --- | --- |
| A166 | T219 | M255 | I292 |
| S169 | F222 | N256 | I296 |
| V170 | V225 | I258 | R297* |
| V172 | E226* | D259** | V299 |
| I173 | T227 | A262 | R300** |
| S176 | C229 | I263 | R303 |
| I177 | I230 | P265 | K306*** |
| S179 | F233 | Y266 | L307 |
| F180 |  | T269 | S308 |
| C181 |  | L270 | R309*** |
| E183* |  | G271 | H310 |
| T184 |  |  |  |

\*salt bridge; \*\*,\*\*\*apparent triads
+other amino acid backbone atoms are included; only side chains are truncated for those amino acids

All side chains that point in, and can interact, and possibly contribute to a proton pathway, are included. 976 atoms stretched the limits of this type of calculation with the computers available to us. An earlier computation included only 672 atoms, truncating more side chains, allowing side chains and some backbone atoms to bulge into vacated space, and producing physically extremely improbable conformations (for one thing, it would be impossible to re-fold these conformations), invalidating the results. The 976 atom case did not allow this; its structure appears to be stable. Voltage made small structural differences, principally to side chains, that turned out to be within the range that could reasonably be expected to return to the initial state with only proton transfers. Given the importance of the residue F233, it is worth noting, for comparison, that six of the seven hydrophobic amino acids discussed in the paper by Schwaiger et al ^30^, emphasizing the role of F233 in gating, are included here, the exception being I231 (We note that Schwaiger et al had the phenylalanine ring rotate after a translation of only 1 Å). The result in our calculation has the phenyl group motion more like a vertical translation of the side chain (i.e., a rotation about a C-C bond), with only a little less rotation than in Schwaiger et al. However, either way the phenyl group with zero volts, hence open channel, is very close to the X-ray structure, while with negative voltage the phenyl group moves in such a way as to prevent a proton coming from below from making contact with the group above. This suggests a H^+^ - phenyl π-electron complex, which should exist, even though it is relatively weak; the complex should not be too strong or the proton could not transfer to the next residue in the chain. It appears that such complexes are of about the right strength for the necessary transfer ^31^. We calculate cases with the protons on the arginines (ionized salt bridge) and on the acid (neutral salt bridge). Both negative (closed state) and zero or positive (open state) fields are imposed on the system. The 976 atom cases each took several months to optimize; running several in parallel made the entire set of computations possible in about one year.

The structures from HF calculations are accurate enough to use; the local energy minima occur in the correct locations to within a fraction of an Å. The errors do not affect interatomic distances by enough to change any conclusions. However, energy values provided by HF calculations may be inaccurate by multiples of k_B_T, even for differences in energy in which most errors cancel. Therefore, single point calculations on the HF optimized structures were carried out at B3LYP/6-31G** level; these required approximately one day each, and are much more accurate. They include the exchange and correlation energy, quantum mechanical effects; the correlation energy is also missing in the HF calculation, and accounts for roughly 1% of the total energy, far greater than the differences in energy that are crucial for the proton transfer. See Table 6 for the values of these quantum contributions to the energy. The differences in the sums of the quantum terms are comparable in magnitude to the total differences between open and closed (*i.e.,* several k_B_T), and they reverse sign when going from open to closed, so that omitting them would produce results with approximately 100% error in the magnitude of the open-closed transition energy.

Energy differences are being compared, so there will be cancellation of systematic error for almost all the system, only a very small part of which changes in single proton steps. With quantum terms included, differences are reliable enough to use in forming a proton path. These energy values are used in the comparison of the structures, in particular in answering the question of whether the proton transfer to a neutral structure is more or less stable than the ionized salt bridge structure. Interatomic distances, for some specific pairs of atoms, show how much, or little, the backbone moved. The distances are compared with those in the X-ray structures for certain key pairs, in Table 3, showing the X-ray structure is closer to the S4 fixed than the S4 free cases (for a more detailed discussion of this point, see Results, part (b), and Discussion, part b).

**Table 2.** Optimized Cases

| S4 | Electric field* | Proton shift** |
| --- | --- | --- |
| Free | Minus | R297 – E183 |
| Free | Minus | No |
| Free | Zero | No |
| Free | Plus | R297 – E183 |
| Fixed | Zero | No |
| Fixed | Minus | No |
| Fixed | Minus | Y266 – E183 |
| Fixed | Minus | R297 – E183 |
| Fixed | Minus | R300 – E183 |
| Fixed | Minus | R297 – E183/Y266 – E226 |
| Fixed | Plus | No |
| Fixed | Plus | R297 – E183 |
| Fixed | Plus | Y266 – E183 |
| Fixed | Plus | R300 – E183 |
| Fixed | Plus | R297 – E183/Y266 – E226 |

*\*the electric field magnitude is $10^7$ V m<sup>-1</sup> along the z-axis, inside negative (closed), or positive (open), corresponding to 70 mV across the membrane; this is the average field, while the local field depends on local charges. If the field is zero, this should reproduce the X-ray structure, presumed to be open. The positive field would correspond to about a potential of +70 mV across the membrane, near the limit of what is possible, but should keep the open conformation.*
*\*\*Proton shifts are from the residue on the left of the pair to the residue on the right. If 2 pairs are shown, two protons were shifted. “No” means that all arginines are positive, all glutamates negative, and Y266 is neutral. For example, R297 – E183 means that the proton shifts from R297 to E183, such that both arginine and glutamate become neutral; if Y266 is shown as the proton source, that residue is negative.*

**TABLE 3.**
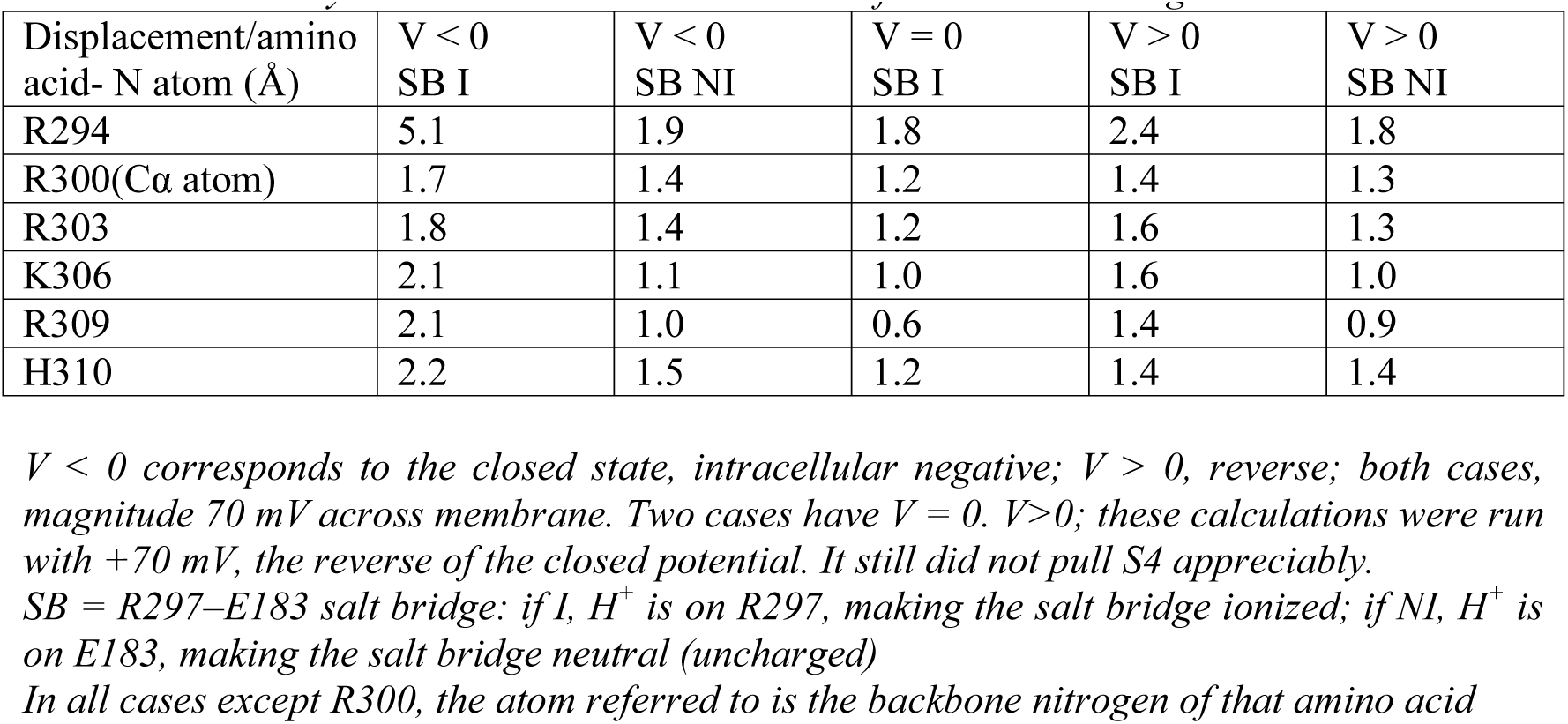
Distances moved by backbone N atoms with various field and salt bridge ionization states

| Displacement/amino acid- N atom (Å) | V < 0<br>SB I | V < 0<br>SB NI | V = 0<br>SB I | V > 0<br>SB I | V > 0<br>SB NI |
| --- | --- | --- | --- | --- | --- |
| R294 | 5.1 | 1.9 | 1.8 | 2.4 | 1.8 |
| R300(Cα atom) | 1.7 | 1.4 | 1.2 | 1.4 | 1.3 |
| R303 | 1.8 | 1.4 | 1.2 | 1.6 | 1.3 |
| K306 | 2.1 | 1.1 | 1.0 | 1.6 | 1.0 |
| R309 | 2.1 | 1.0 | 0.6 | 1.4 | 0.9 |
| H310 | 2.2 | 1.5 | 1.2 | 1.4 | 1.4 |

$V < 0$ corresponds to the closed state, intracellular negative; $V > 0$ , reverse; both cases, magnitude 70 mV across membrane. Two cases have $V = 0$ . $V > 0$ ; these calculations were run with +70 mV, the reverse of the closed potential. It still did not pull S4 appreciably.
SB = R297–E183 salt bridge: if I, $H^+$ is on R297, making the salt bridge ionized; if NI, $H^+$ is on E183, making the salt bridge neutral (uncharged)
In all cases except R300, the atom referred to is the backbone nitrogen of that amino acid

## RESULTS

a. We have calculated the **optimized structures plus energies** for those cases listed in Table 2. A key result is the S4 free, negative voltage (closed state), with all salt bridges ionized; this is the state that the standard models assume must produce a large downward shift of the S4 domain. No such shift is found in these calculations. This is clear from both Fig. 1 and Table 3. Fig. 1 shows the X-ray and optimized structures superimposed, showing that there is almost no motion of S4, except for I292 and R294, which move in the wrong direction. That motion at least shows that the calculation allows motion, if there is any. Table 3 gives quantitative results (part b below). Summary of optimized cases: In all cases the protein starts from the X-ray structure which has protons added by normal mode calculations according to the pdb: 3Lut coordinates. In the cases labeled “proton shift” in Table 2 at least one salt bridge proton from R297 has been transferred to the acid E183, producing a neutralized salt bridge.
b. **Table 3 shows the results of calculations with S4 free to move** against the S1,S2,S3 bundle in terms of distances moved compared to the X-ray structure. The results show essentially no motion. If the field is negative, in standard models the backbone of S4 must be displaced inward to reach the closed state. The results in Table 3 show that this does not happen; there is almost no difference among field negative, positive, or zero. Positive voltage optimizations (equivalent to +70mV across the membrane) were calculated, and are included for completeness. With positive voltage, in principle, the results should be the same as for zero volts, since both should correspond to the same open state. In fact, within a few tenths of an Angstrom, they do agree and both agree within <2 Å with the X-ray structure. When we go to negative voltage, with S4 free, the motion that corresponds to the gating current (intracellular direction, as the channel closes) in standard models is not found; very little motion of any kind is found. The only exception is R294, which moves ≈ 5 Å (and there is some motion of nearby hydrophobic residues) ; while adding nothing to gating current (wrong direction) it does show that when S4 is not constrained, the conditions of the calculation do allow motion. Also, motion in earlier 672 atom calculations was substantial, but into the empty space beyond the edge of the limited system. Because the 672 atom calculations gave results that were obviously erroneous, they are not further discussed here (what motion there was did not agree with any standard model either). With 976 atoms, the structure is maintained. An arginine-phospholipid complex forms, also preventing S4 from moving. (See Discussion, section c). This complex is quite strong, thus providing the physical basis for fixing the S4 end arginines^32^. Some side chains do rotate, affecting local interatomic distances, and thus affecting the probability of proton transfer. We cannot predict that an R294 mutation would necessarily make a huge difference to the channel, as the S4 may also be tied down by another means, such as salt bridge/triad/pentad interactions with S1/S2/S3.Therefore Fig. 1 shows S4 free with negative voltage, compared to the X-ray structure, showing little change in structure, almost none for the backbone; Fig. 2 shows results for the lowest energy structures with no voltage, and with negative voltage, +both with S4 fixed. The shifts in proton position among Y266, R300, and E183 define the low energy cases. Fig. 3 shows the main energy relations among the V=0 and V= -70 mV cases, with the proton shifts labeled.

**Fig. 3:**
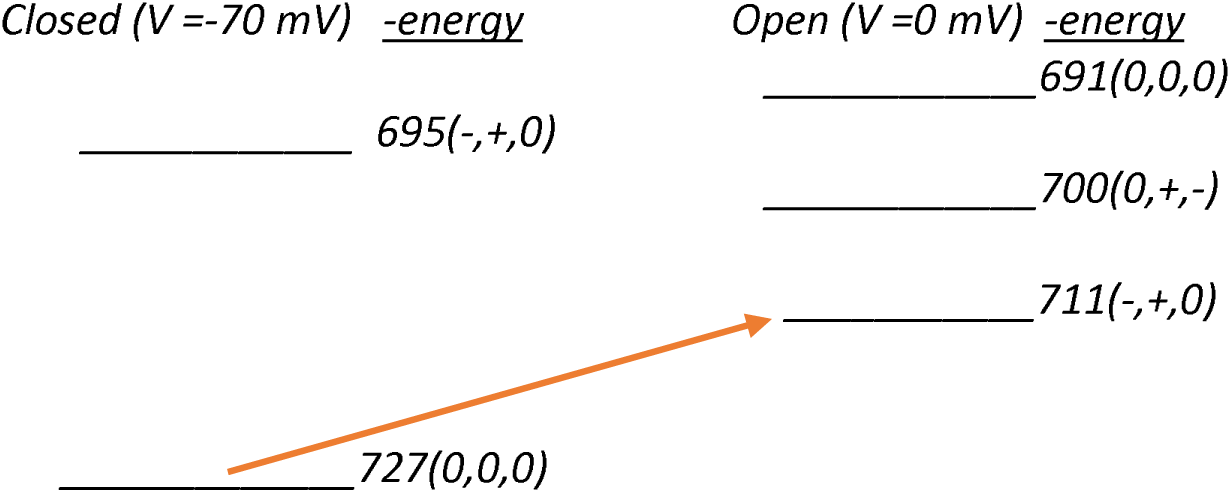
Energy shifts corresponding to proton transitions. As in Table 4, E+25111 Ha. The arrow corresponds to the lowest energy transition between closed and open forms, without accounting for any other intermediate states. The energy difference of about 40 kJ, assuming this is the defining energy, corresponds to a Q_10_ of about 1.7, which is in a reasonable range. How this will fit with other proton transfers (at least one more seems needed) remains for further calculations, but this suggests that most of the energy must come from this transfer.
c. **Proton transitions**: By determining the energy levels for multiple possible proton positions among Y266, R297, and E183, we can get a reasonable estimate of the transition energy for this part of the VSD. Fig. 4 gives the energy values for two closed and three open states (Notation: as in Table 4, (x,y,z)=nominal charge on Y266,R297, E183, respectively, giving the proton locations). The lowest energy in Table 4 is for the closed state, with the proton on the glutamate from R300, making it neutral.

**TABLE 4.**
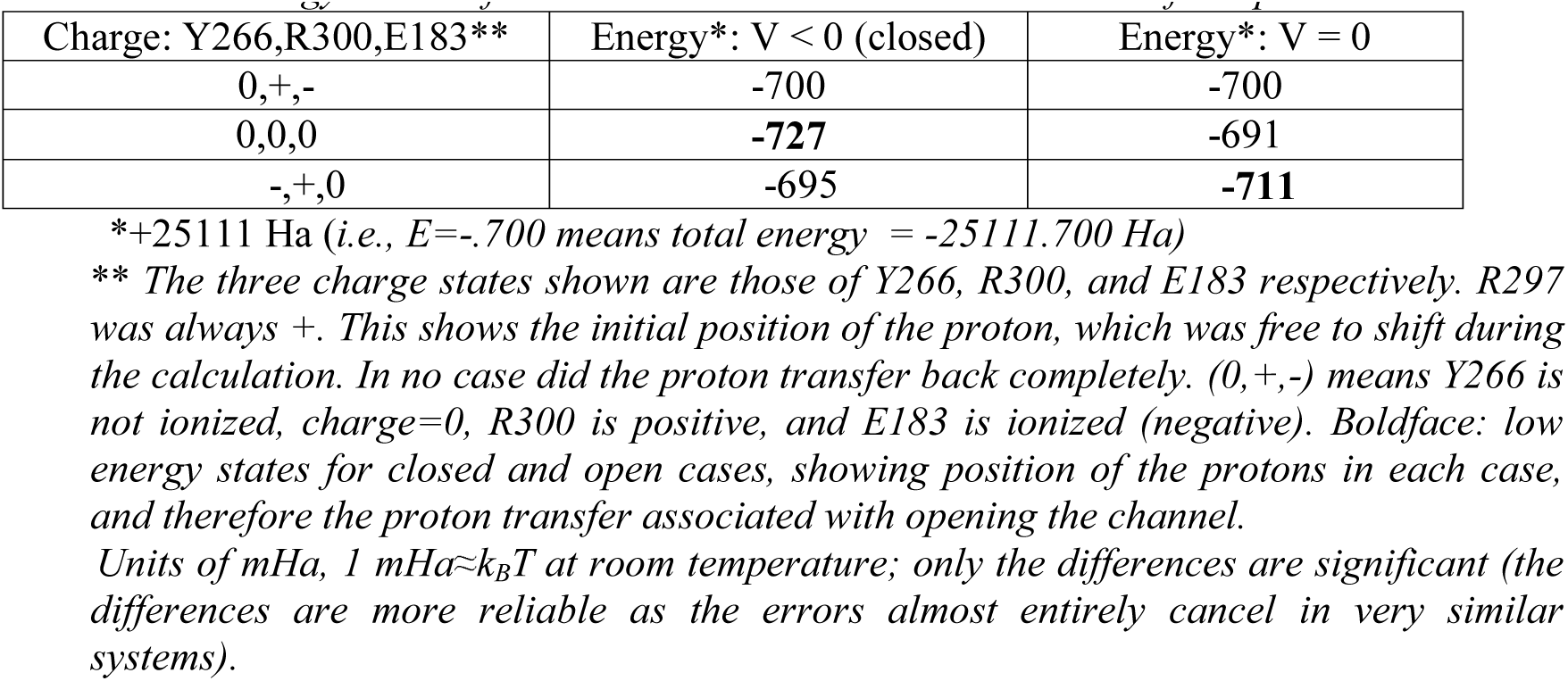
Energy values of cases in which Y266 does/does not transfer a proton

| Charge: Y266,R300,E183** | Energy*: $V < 0$ (closed) | Energy*: $V = 0$ |
| --- | --- | --- |
| 0,+,- | -700 | -700 |
| 0,0,0 | **-727** | -691 |
| -,+,0 | -695 | **-711** |

\*+25111 Ha (i.e., $E = -700$ means total energy = -25111.700 Ha)
\*\* The three charge states shown are those of Y266, R300, and E183 respectively. R297 was always +. This shows the initial position of the proton, which was free to shift during the calculation. In no case did the proton transfer back completely. (0,+,-) means Y266 is not ionized, charge=0, R300 is positive, and E183 is ionized (negative). Boldface: low energy states for closed and open cases, showing position of the protons in each case, and therefore the proton transfer associated with opening the channel.
Units of mHa, $1 \text{ mHa} \approx k_B T$ at room temperature; only the differences are significant (the differences are more reliable as the errors almost entirely cancel in very similar systems).

**Fig. 4:**
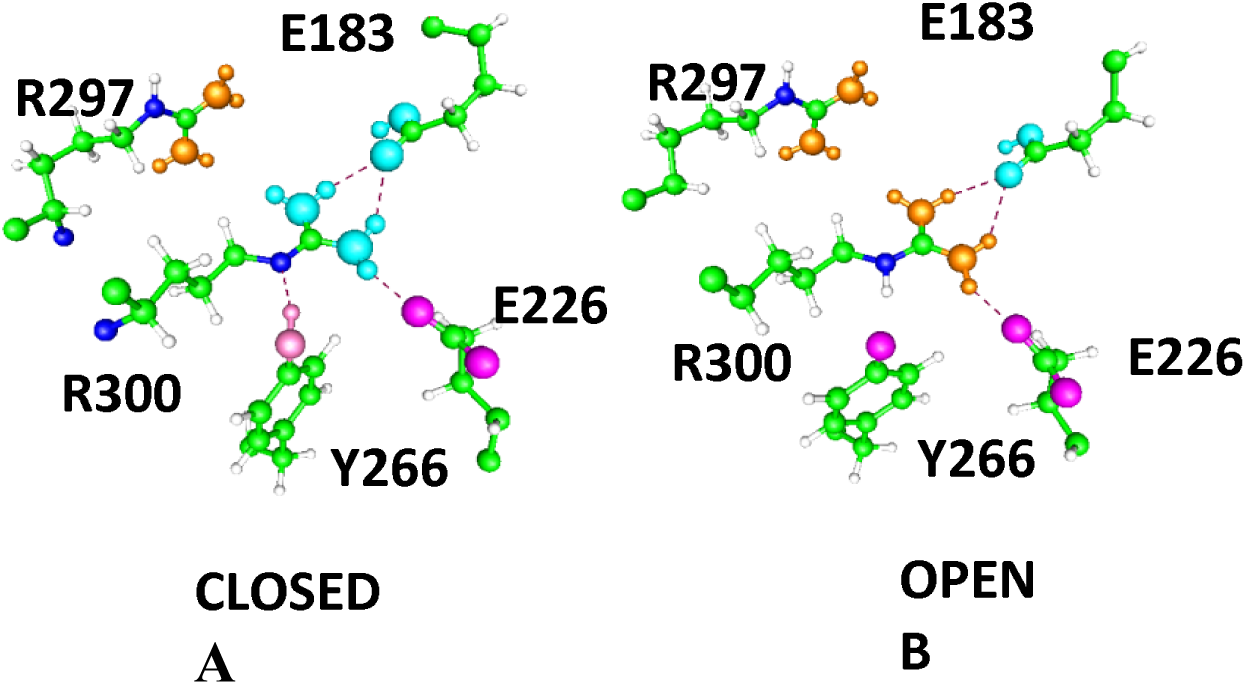
A key proton transfer involving Y266; the proton moves from the tyrosine in the closed state to the nitrogen on the R300 in the open state. The guanidinium group is shown as blue when R300 is neutral (A) and gold when it is charged (B). The shift also influences the position of the side chain of E183, so that a two step shift (Y266➔E183) can occur, with R300 serving as pathway; however, E183 appears to be neutral in both open and closed states. (For energies, see Fig. 3 above). In addition, when the proton is on R297 (as shown for open) the Y266 sidechain rotates downward toward R303 at almost the same energy, so that the position of Y266 is not as well defined as in the closed state, where Y266 is locked, in the position shown in the figure above. The lowest energy open state, is approximately 16 k_B_T above this state, suggesting that for this state at least there is about a 40 kJ activation energy. This is in line with a number of measurements of Q10, and corresponds to a Q10 of approximately 1.5, somewhere in the middle of the range of experimental values.
d. **Figs. 3 and 4 illustrate a key shift by the protons**. See also section (h), below. Several other transfers were calculated, but led to transitions to states that were higher in energy.
e. **Salt bridge ionization:** The fact that some of the salt bridges need not be ionized is not a surprise: water is required to ionize a salt bridge (other polar solvents are not relevant here). One water is not enough, two may be sufficient, and three certainly are ^13a^. In the center of the VSD, there is a hydrophobic region in which there is no water, and we do not expect ionization, so shifting a proton to neutralize the two components of the salt bridge at that point is expected. In the R➔C mutant, however, there is room for two or three water molecules, so the ionization state could easily be different; the entire channel should behave somewhat differently. Furthermore, it is possible to find structures that suggest the type of proton resonance that appears in much simpler calculations of isolated salt bridges with 2 or 3 water molecules^13b^. Such structures help to stabilize the charge transfer, and the seemingly anomalous pK values. These are quantum effects, and would not appear with any form of classical potentials, as far as we can determine.
f. **Charges**: Table 5 gives atomic charges under various conditions. Charges on key groups: the arginines, glutamates, and tyrosine have non-integral charges; the electron wave functions produce a charge distribution among the atoms that shows the fraction of charge on the various groups. There is shared charge.

**Table 5a.** Charges on Y266 and glutamates*

|  | 0 field; R300(+) | V>0, R300(+) | V>0, R300(0) | V<0, R300(+) | V<0, R300(0) |
| --- | --- | --- | --- | --- | --- |
| Y266 | -0.241 | -0.843 | -0.247 | -0.843 | -0.248 |
| E183 | -0.742 | -0.014 | -0.016 | -0.009 | -0.013 |
| E226 | -0.720 | -0.733 | -0.729 | -0.732 | -0.728 |

\*For Y266, the charge is either on the O if ionized, (so R300(+)), or else if Y266 is not ionized (so R300(0)); for glutamates, charge is the sum of that on the carboxyl oxygens plus the carbon to which they are bound, plus the proton if the carboxyl includes it (not ionized case, which applies to all except the first column; the E183 proton comes from Y266 in columns 2 and 4, and from R300 in columns 3 and 5).

**TABLE 5b.** Charges on arginines* We see first that the charges are not particularly close to integral, meaning that the electrons are spread out over several atoms, and the charges likewise. This is not a surprise, as comparison with a particle in a box would suggest that a bigger box means a lower energy. Concentrating charge should increase electrostatic repulsion. As a result, we ought to expect *a priori* a result like the one that is found, rather than integral charge on groups. This does have implications for proton transfer, as the electronic wave function forms a partial bridge between groups. There may be some effect of delocalization error^33^ (charges are calculated with B3LYP, a DFT method), but this will not be large enough to affect the overall result. It is not required that the path followed by the protons is the same for the open➔closed and the closed➔open transitions. Instead, it appears more probable, based on the charges that must exist at intermediate states, combined with the field, that two different paths are required. It is known that there is hysteresis in the gating transitions ^26^. The existence of multiple states from closed to open has been suggested on the basis of kinetic data on channel opening by many workers, which supports the idea that more than one path may exist.
g. **Role for Y266**: The shifts in pK values that appear to exist, based on these calculations, are not simply a consequence of local electric fields. We have been discussing Y266, which, in the open state, moves so as to place its −OH group near an R300 nitrogen of the guanidinium that has lost its proton. The hydrogen from the Y266 −OH group is partially shared with this nitrogen, making the R300 deprotonation incomplete, and allowing a short chain of partially bonded protons to form. Thus the R300 deprotonation does not require so large a pK shift as it would in solution. Fig. 3 and Table 5 (actual charges) show how this affects energy and charge, and Fig. 4 shows the local structural relation. This step is itself an intermediate step in gating. We have calculated 13 cases with S4 fixed, (4 with S4 free). States with high energy are not discussed. In the following, the nominal charge is in parentheses, corresponding to proton position. The overall system is unchanged except for the proton transfers. The summary of these steps is that a charge has moved from Y266 up to E183 upon. In the closed state the ionized Y266 corresponds to high energy, and in the open state to low energy, so the proton must transfer in opening the channel. The transfer is in the correct direction (proton moves upward). In Fig. 4, the step shown corresponds to transfer of the proton from Y266 (closed) to R300 (open). This is a step that was calculated, but another step (Fig. 3) shows the energies of relevant open and closed states, and thus shows the lowest energy states.
h. **Charge shifts on opening the channel:** Combining the results shown in Table 5 with those in the preceding paragraph, we have the change in charge on the relevant residues on opening the channel. This shows approximately 0.5-0.6 charges having transferred, in moving from Y266 to R300, so that this step provides about 20-25% of the gating current for the entire VSD. This is not unreasonable for a single (set of) step(s). It is consistent with the need for at least one more H^+^ transfer, and a possible contribution from the dipoles, but these latter transitions remain for future calculation.
i. **Role for F233**: Several side chains were fairly mobile. F233 is particularly important, folding downward to follow the proton when it transfers (Fig. 2). This dihedral angle rotation, accompanying the proton shifts, appears to be a significant difference between open and closed states. F233 is well conserved, and appears to be central to the proton transfer mechanism. A similar phenylalanine exists in H_v_1, and proton transfer control in H_v_1 has been attributed, at least in part, to its dihedral angle transition ^21a^. It appears to provide a hydrophobic separation between two proton paths, but may connect the protons with an aromatic π electron to H^+^ complex.
j. **Exchange and Correlation Energy** The differences between V= 0 mV and V = -70 mV are significant, the sums (exchange + correlation) being different (ionized(0mV) − unionized(0mV)) for open channel by 28 k_B_T, and by 3 k_B_T (two closed states, favoring unionized). The relaxed form (0 mv, unionized) has the lowest exchange plus correlation energy, and presumably corresponds to the normal open state. These differences are comparable, or even larger than, the total energy differences, so that ignoring the quantum contributions to the energy would lead to approximately 100% error in the energy differences.
k. **Tests of the accuracy of the calculation**: There are two tests, one offering comparison with experiment, the other a plausibility argument. **1)** structures, especially key interatomic distances, in the proton-shifted (neutral salt bridge) cases, with zero voltage, S4 fixed, came out close to the X-ray structure, as they must. This also suggests that the structures resulting from these calculations are realistic. The S4 free structures are not quite so close. Had the S4 free structures come out closer, we would have concluded that S4 free was the correct structure. If neither had, we would have concluded that the calculation itself was at fault. However, the fact that the structures that gave the closest replica of the X-ray structures were S4 fixed, field = 0, and were quite close, the results strongly suggests that the calculation is correct, and the S4 fixed structures can be trusted to within a fraction of an Angstrom, especially for key salt bridges and bond lengths. By extension, we trust the negative voltage results, although no X-ray structures exist. Other data support this assumption. The key R297 − E183 salt bridge, with applied negative potential, which should have broken if S4 were to move down in order to reach a closed position, instead tightened from approximately 6 Å (guanidinium N to carboxyl O) to approximately 4 Å, when the salt bridge was charged, with S4 free, as assumed in standard models. For the salt bridge not charged (*i.e.,* H^+^ transferred from arginine to glutamate), there should be almost no response to the field (only a dipole remains, and the interaction energy with the field would be too small to show any structural effect); nothing in the calculation contradicts this point, and the same result would be expected on any model. However, the uncharged closed case has lower energy, strongly suggesting that the proton has transferred from arginine to glutamate. Standard models that require physical motion at constant charge to open the channel are not consistent with this result. **2)** The difference in energy of the salt bridge ionized and unionized structures is generally of the order of 10 k_B_T, often less than 10 k_B_T, making it possible for the charge to shift back and forth with the field. The one large barrier is the energy difference between ionized and unionized Y266 in open and closed states, where a large barrier appears to lock down the closed state, while the reversal of this barrier allows the proton transfer required for the channel to open. The calculation suggests that the proton has already continued on to E183 in the closed state, with the opening transfer only from Y266 to R297. This implies that the gating charge accompanying this step is fairly close to 0.5 − 0.6 as suggested earlier. While only a plausibility argument, an erroneous calculation would be certainly expected to produce much larger differences in energy, as the independent calculations would then fail to be closely related.
l. **Dipoles**: We earlier discussed the dipole of the water group alone. However, the dipole of the entire system leads to another question that must be considered. Table 7 gives the total dipole for the two low energy states, plus the fully ionized model that is assumed in all standard models. The dipole moments on the left correspond to the case of the closed channel, energy (+25111)=-.727, the middle case is the open channel, E=-.711, and the channel as assumed in standard models, with all H on arginine, and tyrosine neutral, has E =-.700, channel open but about 11k_B_T higher energy than the proton shift shown. The changes in the dipole moment might contribute to the classical part of the energy, but the total **μ*E** term, taking everything as homogeneous, is an order of magnitude too small to account for the effects observed. If the field is sufficiently inhomogeneous, and the dipole is large where the field is large, the contribution may become substantial. We cannot yet say that that can be ruled out. If that is the case, the rotation of the dipole, not evident in the overall value shown here, could contribute substantially to the gating current. The water dipole (Fig. 5, Table 8) is about 2/3 of **μ**_X_, and does rotate, in the calculation; more work is needed to understand the real system. **Water dipoles** The dipole moments for these droplets are given in Table 8, showing the rotation of the dipole from parallel to field, meaning approximately parallel to channel axis (closed, with field) to orthogonal to the axis (open, no field). The droplet shown in Fig. 5 consists of the water molecules as optimized in the complete 976 atom system; those coordinates are used for a single point B3LYP/6-31 G++ calculation of energy and dipole of the water molecules alone. We cannot draw quantitative conclusions from the water droplet; the number of water molecules is approximately what fit in the space with the VSD, hence reasonable, but the exact number is affected by lipids, and the neighboring solution, both omitted from the calculation. These water molecules form a reasonable boundary between the VSD and the missing external neighboring molecules in the calculation, at the high-dielectric extracellular end. (We draw conclusions primarily from the interior of the remainder of the protein; the vacuum boundary in the remaining region would be relatively low dielectric constant in the complete system. The vacuum boundary limits the accuracy of the calculation for some groups near the boundary, like R294). Table 8 shows the rotation of the dipole from parallel to field, meaning approximately parallel to channel axis (closed, with field) to orthogonal to the axis (open, no field). The majority of the total water dipole rotates from alignment along the X coordinate, through the VSD, when the field is off, to mostly orthogonal to that direction, along the Z coordinate parallel to the plane of the membrane, with field negative (closed). The difference would correspond to an appreciable fraction of the gating charge, but because it cannot be taken as quantitative, we only note that the water could make a significant contribution to the gating charge. The total dipole of the system, which is only about 70% greater than the water dipole does not appear to rotate like the water dipole, and there is good reason to not assume that the dipole is even approximately homogeneous through the system. So far we cannot estimate the contribution of rotating dipoles to the gating current; it may be an appreciable fraction of the entire gating current, but we cannot yet say how large a fraction.

|  | 0 field;<br>R300(+) | V>0, R300(+) | V>0, R300(0) | V<0, R300(+) | V<0, R300(0) |
| --- | --- | --- | --- | --- | --- |
| R297(6) | 0.144 | 0.158 | 0.162 | 0.147 | 0.160 |
| R297(7) | 0.829 | 0.838 | 0.842 | 0.835 | 0.840 |
| R297(9) | 0.635 | 0.651 | 0.653 | 0.655 | 0.653 |
| R300(6) | 0.126 | 0.091 | 0.070 | 0.093 | -0.071 |
| R300(7) | 0.813 | 0.778 | 0.572 | 0.781 | 0.570 |
| R300(9) | 0.620 | 0.622 | -0.094 | 0.624 | -0.095 |
| R303(6) | 0.142 | 0.116 | 0.135 | 0.111 | 0.134 |
| R303(7) | 0.838 | 0.798 | 0.817 | 0.793 | 0.816 |
| R303(9) | 0.627 | 0.577 | 0.599 | 0.577 | 0.598 |

\*Charges on each arginine are summed over 6, 7, or 9 atoms: 6 sums over both $\text{NH}_2$ groups, or $\text{NH}+\text{NH}_2$ if one proton has transferred to another group; 7, add the central carbon; 9, add the preceding (nearer backbone) $\text{NH}$ . i.e., if the $\text{H}^+$ from arginine has transferred to another group, the sum over two $\text{NH}_2$ groups becomes the sum over $\text{NH} + \text{NH}_2$ and the number of atoms becomes 5, 6, or 8.

**Fig. 5:**
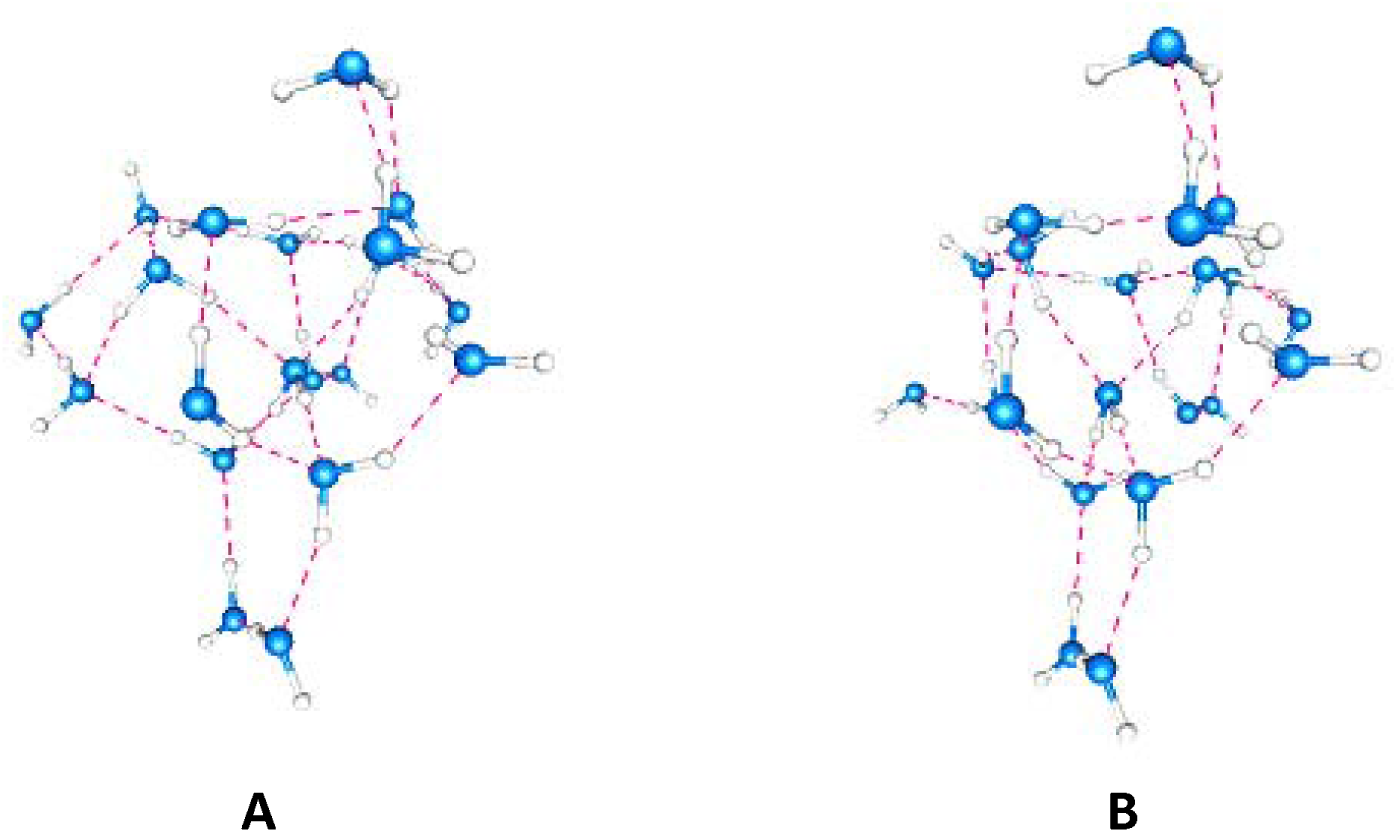
Water cluster for open (A) and closed (B) cases.

**TABLE 6.**
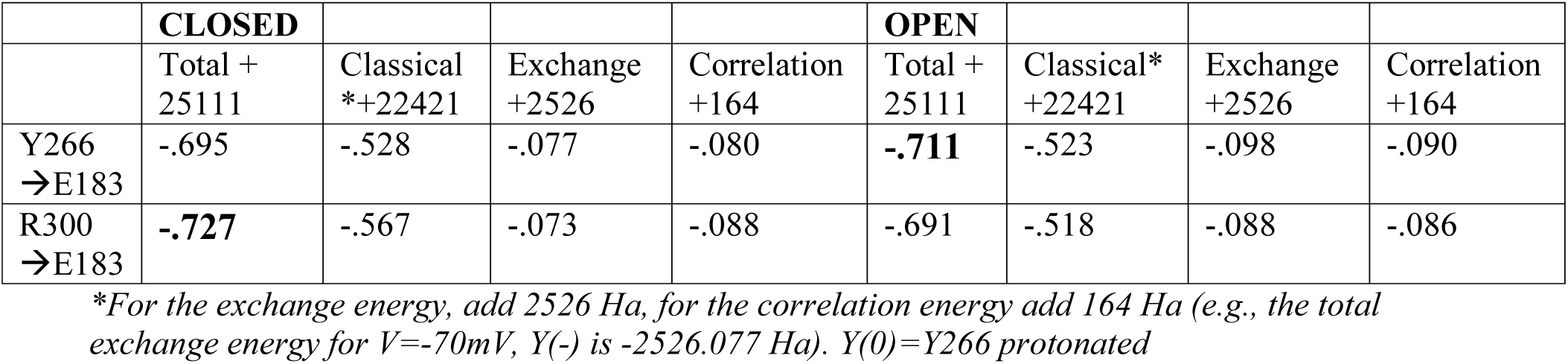
CLASSICAL, EXCHANGE AND CORRELATION ENERGY TERMS FOR THE TWO LOW ENERGY CASES

**TABLE 7:** DIPOLE MOMENTS OF THREE LOW ENERGY STATES+

| V<0 |  | V=0 |  | V=0 |  |
| --- | --- | --- | --- | --- | --- |
| H shift:<br>R300→E183 |  | H shift:<br>Y266→E183 |  | No H shift, salt<br>bridges ionized |  |
| $\mu_x$ | $\mu^*$ | $\mu_x$ | $\mu^*$ | $\mu_x$ | $\mu^*$ |
| 27.1 | 31.6 | 24.8 | 28.0 | 52.3 | 56.3 |

<sup>+</sup> $\mu_x$ is the component along the channel axis, $\mu^*$ the total dipole of the 976 atom system

**TABLE 8.** Properties of the water cluster: energy and dipole moment

|  | Energy** | Total dipole (D) | X component | Y component | Z component |
| --- | --- | --- | --- | --- | --- |
| Zero field | -.203 | 16.9 | 16.0 | 5.1 | -1.4 |
| $-10^7 \text{ Vm}^{-1}$ along X* | -.199 | 17.4 | -1.3 | 4.7 | 16.7 |

\*Field negative (closed), corresponding to $-70 \text{ mV}$ along X. 1 Debye (D) is $3.33 \times 10^{-30} \text{ C m}$ .
\*\*Hartrees + 1605. (i.e. total energy -1605.203, -1605.199; closed conformation has about $4k_B T$ higher energy, with $1 \text{ mHa} \approx 1 k_B T$ .)

## DISCUSSION

The results point to a mechanism for gating current that depends on the shift of protons rather than a shift of the physical location of the backbone. We can summarize what we have found so far at this point, with the addition of consideration of the implications for a gating mechanism. Some parts are only a hypothesis, but are required as part of a complete coherent mechanism.

## a) Gating steps, with a hypothesis

So far, we have stated that A) the gating current is carried primarily by protons instead of S4 motion, possibly with a contribution from water at the extracellular end, and B) the ionization state of the salt bridges can be influenced by neighboring residues, allowing proton transfer. We must understand B in order to understand A; furthermore, we extend “salt bridges” to include larger groupings of amino acids that can join in transferring a proton. Therefore, let us look in some detail at certain triads of amino acids that are encountered in the VSD. Similar triads are found in cyt-c and in H_v_1. The existence of these triads suggests a path for the protons to follow as their transfers create a gating current. However, our calculations show that the salt bridges are not merely influenced by third amino acids, but that there are transfers of a proton through an intermediate residue between two neighboring sets of residues; the final position of the proton allows it to move to the next grouping of three or more amino acids. Fig. 4 shows a grouping in which a single transfer is influenced by its neighbors. A single residue can participate in two neighboring triads; alternatively, the grouping can be considered a single grouping of five. An initial ionization/proton transfer shifts the local potential so that the next transfer in the sequence occurs, and the remainder of the shifts that generate gating current follow. The initial pulse of gating current (the “piquito”) ^17^ is too fast to measure, and seems consistent, in speed, and size of the effect, with proton tunneling^11, 15^ although an alternate possibility has been suggested^17a^. We take the “piquito” to be, with high probability, a proton tunneling event; we cannot yet state which amino acids constitute the key pair. However, there is a large field drop, apparently constituting most of the voltage drop across the membrane, near R297, so the transfer of a proton there is a candidate for the possible key initial step ^34^.

Triads may be composed of two bases and an acid, two acids and a base, or may have an alcohol, here Y266, in place of one of these; hydrophobic amino acids are not included. In principle, with the aid of tautomerism, even the amides (glutamine, asparagine) might participate, but we do not find a case of this here. The base is not limited to arginine or lysine, but may also be histidine. The proton path in the VSD appears to be surrounded by hydrophobic amino acids, which may help define the structure as well as keep water out, so that the random orientation of a water molecule cannot be a problem. It is not difficult to see how a proton could transfer along the acid/base/alcohol path, with the assistance of a change in external electric field. Our proposed proton gating path for the K_v_1.2 channel includes the following triads: 1) E183, R297, Y266, R300; 2) R303, E226, R300. These two overlap at R300, making a pathway that goes through the center of the VSD; taken together these can be regarded as a single sextet of amino acids. Because these overlap, they can form alternate paths. Interrupting either one separately does not kill the channel although it will change the gating current somewhat. Everything so far is included in the calculations already completed. We have not yet found an alternate path for this section, but expect that such a path could be found. Because F233 is directly below R300, and the π electrons of the phenylalanine can deflect a proton, further computations with this residue are likely to be needed. The electric field interacts with the gating charge by moving the protons, which become delocalized.

The remainder of this section constitutes a hypothesis, with computations on these sections just beginning: The H^+^ on opening comes from (R303,E226), with the path beginning at the lower end of the VSD: the path includes 4) K306,D259,R309,R240,E236; 5) extends through S169,H310, and 2 water molecules. The step from E226, R303 to K306, D259 is short enough that there should not be a major barrier (the role of F233 remains to be determined in this regard, however.). The system continues on toward H418, which is hypothesized to have an added proton in the closed state, and the conserved PVPV sequence at the gate. The proton, on moving up, allows the gate to open. Both classical and quantum terms depend on the location of the proton. The PVPV sequences of the four domains are spaced far enough apart that the actual gate appears to be at the junction of the PVPV sequence with the upper part of the T1 moiety that hangs below it, where the opening is smaller. Since it is sometimes possible to get a functioning channel with T1 truncated, it may be that a path through the water at the inner surface of the membrane can organize to transmit a proton, although exactly how the gate closes is not yet clear’ a move of 2 − 3 Å should be sufficient. The central part of the K_v_1.2 sequence is similar to H_v_1; the R300-E226- R303 triad for K_v_1.2, is comparable to R201, D108, R204 for H_v_1: the distance of the closest N atom of R303 to the nearest carboxyl of E226 is 4.26 Å (X-ray distance), the corresponding distance for H_v_1 is 4.34 (R201 – D108); corresponding distances for R300 to E226 = 2.71 Å (K_v_1.2) and R204 to E108 = 2.71 Å also (H_v_1). The two channels diverge below this. The H_v_1 proton pathway continues toward the cytoplasmic side through several sets of residues that, taken together, produce a pathway that is somewhat similar to that in the VSD of K_v_1.2, albeit differently oriented, and with different residues. All this said, everything in this paragraph requires further computation.

## b) S4 free vs S4 fixed

Section (b) of the Results section gave reasons based on the calculations for taking S4 fixed. In addition, there is a physical mechanism for keeping it fixed; it is likely that the end of the S4 segment is held by the membrane. The first arginine (R294) is in the membrane region, near phosphate groups. At least some negative charges must exist in the membrane to have a functioning channel^35^, and the mechanical state of the membrane also matters ^36^; this seems unlikely if S4 simply went through it. Arginines form fairly strong complexes with phosphates ^32, 37^, which would anchor R294, making a fixed position of S4 unavoidable. If such a complex were to form, it would have to be pushed apart by the force on the S4 breaking out of the central part of the membrane, which appears to require far more energy than is available. This is further evidence that S4 fixed cases appear to be the correct ones for the interpretation of the results, and for understanding the gating current. Table 3 shows how small the motions are of the S4 segment with the S4 free, V<0.

## c) The role of Y266

We have several times mentioned that Y266 is a key residue in the sequence of proton transfer steps. This residue can provide a proton to R300, which in turn passes it to E183 and R297. R300 is oriented slightly differently than the other arginines so that it allows the transfer of a proton from Y266. The proton donation from Y266 to R300 is one of two paths that can provide gating charge (section b above). A deprotonated tyrosine also can act as a valve to insure unidirectional proton transfer in the K-channel analog of B-type cyt-c oxidase^23^. That system has a somewhat different chain than the one we observe in the VSD, but still uses a tyrosine as a valve for proton transfer. Another voltage sensor based on tyrosine has recently been reported for the M2 muscarinic receptor^38^, albeit with a different interpretation. Our calculation predicts that Y266 mutations would make an appreciable change in gating.

## d) Charges on several amino acids (see Table 5)

The NBO charge calculations give us a more accurate value for the charge transfer accompanying proton shifts. In some cases the proton final position is in the center of a short bond, and it is important to know where the charge resides. Charges are assigned to each atom by the calculation. The charges are not close to integral for the guanidinium or for the acid. It is too simple to say that the charge has transferred as a unit; partial charge transfer, with electron density shared among atoms, and groups, is found. This must be taken into account in any calculations done on the system.

## e) Our hypothesis agrees with previous experimental results on H418

The final (hypothesized) steps include protonation of H418, which is close to the gate at the C-terminal end of the channel, and is expected to play a key role in opening and closing the channel. Deleting this residue has been reported to kill the channel^39^.

We conclude that the gating mechanism need not involve motion of an entire transmembrane segment of protein, and that the gating current, as well as the VSD/gate interaction, can be accounted for by motion of H^+^ through the VSD, possibly with some contribution by the water dipoles at the extracellular end of the channel.

## Summary

In this paper, we present quantum calculations that include a large fraction of the VSD. Our results show a gating current composed, at least in large part, of proton motion, with an essentially immobile S4 transmembrane segment. Quantum calculations are generally limited by the size of the system that can be calculated. We here present a set of calculations larger, to the best of our knowledge, than any previously offered. We believe these to be large enough to adequately represent much of the VSD, and they include an electric field applied to the VSD. Previous work has generally been understood in terms of gating models with mobile S4 segments of the VSD providing the gating current. In earlier reviews, we have considered reinterpretation of the evidence that has hitherto been used to support the idea of S4 motion, particularly SCAM results and MD calculations ^11b, 40^; we showed that this evidence could be understood in terms of the essentially static S4 segment we see in our present results. Our new calculations enable us to propose specific proton transfers in part of the channel, and hypothesize additional transfers that would continue the chain, thus providing the gating current and opening and closing the channel. Further work is needed to complete the proton transfer chain, so as to fully understand gating in the channel.

## Author contributions

A. M. Kariev and M. E. Green designed and carried out research, and analyzed results. M.E. Green wrote the paper

## ACKNOWLEDGMENTS

This research was supported, in part, by a grant of computer time from the City University of New York High Performance Computing Center under NSF Grants CNS-0855217, CNS-0958379 and ACI-1126113, and this research used resources of the Center for Functional Nanomaterials, which is a U.S. DOE Office of Science Facility, at Brookhaven National Laboratory under Contract No. DE-SC0012704.

